# High genetic diversification in a symbiotic marine annelid is driven by microgeography and glaciation

**DOI:** 10.1101/2023.10.13.562162

**Authors:** Yui Sato, Laetitia G.E. Wilkins, Alexander Gruhl, Harald Gruber-Vodicka, Nicole Dubilier

**Affiliations:** Max Planck Institute for Marine Microbiology, Celsiusstr. 1 Bremen 28359, Germany

**Keywords:** biodiversity, speciation, microevolution, genetic connectivity, microgeographic population structure, last glacial maximum, meiofauna, low-coverage sequencing, in-silico exome capturing, gutless oligochaete

## Abstract

Marine invertebrates with limited dispersal abilities exhibit high levels of genetic divergence among populations. However, the spatial extent of genetic differentiation in these species remains poorly understood because identifying natural barriers to gene flow can be challenging in the marine environment. In this study, we investigated the population genetic structure of the interstitial annelid *Olavius algarvensis*, a species that lays eggs in its immediate surroundings and does not have an active dispersal phase. We analyzed the mitochondrial and nuclear genome sequences of hundreds to thousands of individuals from eleven sites in the Mediterranean, spanning microgeographic scales of < 5 km to macrogeographic scales of 800 km. Comparisons of single nucleotide polymorphisms (SNPs) in mitochondrial genomes revealed a complex history of introgression events, with as many as six mitochondrial lineages co-occurring in individuals from the same site. In contrast, SNP analyses of nuclear genomes revealed clear genetic differentiation at micro- and macrographic scales, characterised by a significant isolation by distance pattern (IBD). IBD patterns further indicated the presence of a historical physical barrier to gene flow on the east coast of the island of Elba corresponding to the historical shoreline around Elba during the Last Glacial Maximum in the Late Pleistocene, and highlighting the influence of geological forces in shaping population genetic structuring in the species today. Overall, our results provide strong empirical evidence for the high genomic diversification across spatial scales in marine interstitial fauna.

## Introduction

Understanding how adaptation and diversification lead to speciation requires characterizing the spatial scale of population genetic divergence (Palumbi 1994; Presgraves 2010). Divergence among populations is driven by genetic drift and local selection, but gene flow through dispersal of genotypes can swamp genetic differentiation among populations (Yeaman and Whitlock 2011; Savolainen et al. 2013; Sexton et al. 2014). Therefore, species with limited dispersal abilities are expected to exhibit high levels of population structuring at small spatial scales, increasing the likelihood of genomic diversification and speciation potential (Bradbury et al. 2008; Pelc et al. 2009). However, the geographic extent of genetic divergence remains poorly understood in marine species because defining barries to their gene flow is often challenging (Kelly and Palumbi 2010; Cerca et al. 2018). This is particularly difficult for small, cryptic marine invertebrates.

Microscopic invertebrates dwelling in marine sediments, commonly referred to as the meiofauna or meiobenthos, consist of a diverse range of animals including nematodes, copepods, flatworms, ciliates, annelids, and many other animal groups (Giere 2009). These interstitial animals have raised extensive debates in the field of biogeography because many of the individual species occur over large geographic regions despite their short lifespans and the lack of obvious dispersal mechanisms (Giere 2009; Cerca et al. 2018). This so-called ‘meiofauna cosmopolitan paradox’ has prompted studies of dispersal capacity and phylogeographic patterns of meiofaunal taxa (Todaro et al. 1996; Westheide and Schmidt 2003; Curini-Galletti et al. 2012; Narjes et al. 2017; Struck and Cerca 2019; Worsaae et al. 2019). Experiments of passive and active dispersal under controlled conditions provided evidence for dispersal potentials in some meiofaunal species, e.g., nematodes, rotifers, tardigrades, ostracods, copepods and foraminifera, suggesting possible explanations for their apparent cosmopolitan patterns (Hagerman and Rieger 1981; Chandler and Fleeger 1983; Palmer 1986; Arroyo et al. 2006; Radziejewska et al. 2006; Fontaneto 2019). Accordingly, the phylogeography of marker genes suggested broad distributions and historic events of regional migration in some meiofaunal species such as nematodes and annelids (Derycke et al. 2008; Narjes et al. 2017; Worsaae et al. 2019). However, approaches using maker genes often lack sufficient resolution required to understand population structuring across smaller spatiotemporal scales. For example in gastrotrichs and annelids, counter evidence against the meiofauna cosmopolitan paradox has accumulated from genomic fingerprinting and morphological ultrastructure analyses, indicating that distinct sub-species groups do exist across oceanic regions (Todaro et al. 1996; Schmidt and Westheide 2000; Giere 2009; Zakas and Wares 2012; Kieneke and Nikoukar 2017; Cerca et al. 2018). To elucidate the genomic diversification in interstitial marine invertebrates further, it is necessary to study their population structure across spatial scales from local to regional. This requires population-wide whole genome sequencing to achieve sufficient genetic resolution. Recent advances in whole genome sequencing technologies and Bayesian approaches for identifying single nucleotide polymorphisms (SNPs) have made such population genomic studies possible in non-model species (Andrews et al. 2016; Fuentes-Pardo and Ruzzante 2017). Here we studied the population structuring of a marine interstitial annelid at a microgeographic scale by applying an exomewide SNP-identification method that is based on low-coverage sequencing and posterior genotype probabilities (Therkildsen and Palumbi 2016).

*Olavius algarvensis* occurs in sandy sediments around seagrass beds in the Mediterranean region and often represents the most abundant member of the interstitial fauna (Giere et al. 1998; Giere and Erséus 2002). Their bright white body reaching a size up to 4 cm in length and 200 µm in diameter makes it easy to collect them from the sediments (Figure 1a). Most importantly, *O. algarvensis* has a restricted dispersal potential because the oviposition and juvenile development occur within the sediment, and pelagic migratory stages are absent (Giere and Erséus 2002; Giere 2006). These characteristics make *O. algarvensis* an ideal subject for assessing the potential of population genetic diversification at microgeographic scales.

**Figure 1.**
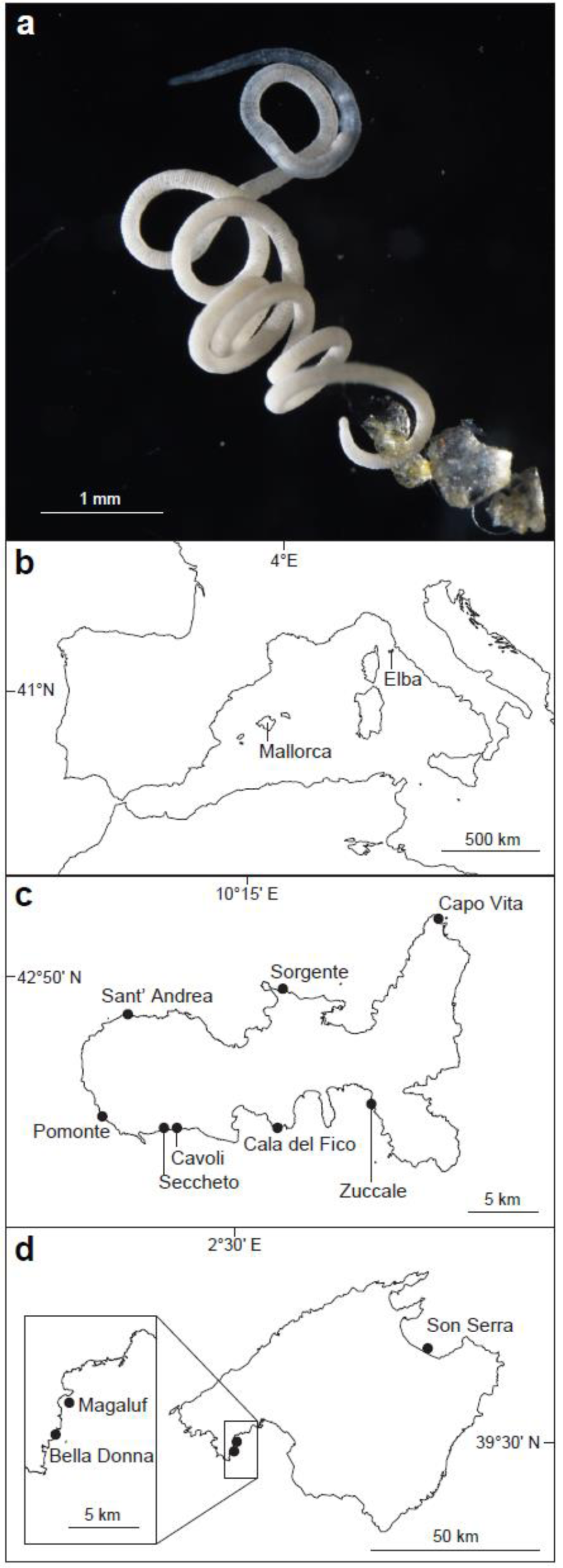
The marine gutless annelid *Olavius algarvensis* and the geographic locations of the study sites. (**a)** Microscopy image of *O. algarvensis* **(b)** Geographic locations of Mallorca and Elba islands **(c)** Study sites around Elba **(d)** Study sites around Mallorca.

To characterise the population structuring of *O. algarvensis* at a microgeographic scale, we compared genomes of individual animals collected in eight bays around the small island of Elba across a 60 km stretch of the coastline (Figure 1b-c). The nuclear genome divergence between *O. algarvensis* populations was assessed among locations based on exomewide SNPs. Nuclear genomes indicate the contemporary genomic connectivity and reproductive isolation among populations, as genetic recombination continually mixes genomic composition among individuals unless they are reproductively isolated. The phylogeny of the mitochondrial genomes was also examined to compare nuclear genetic divergence with the evolutionary history of mitochondrial lineages. Unlike nuclear genomes, mitochondrial lineages are inherited vertically (mostly maternally) over many generations within a population (Hellberg et al. 2002). Therefore their phylogeography can reveal historical demographic introgression events (e.g., Schmidt et al. 2017; Thanou et al. 2019; Mackiewicz et al. 2022). A previous study of *O. algarvensis* in two bays around Elba indicated that multiple divergent mitochondrial lineages co-exist within each bay, suggesting a complex demographic history (Sato et al. 2022). However, nuclear genomes of these mitochondrial lineages have not been investigated and thus the pattern of reproductive isolation of *O. algarvensis* around the Elba locations is unknown. To extend our insights into population genetic diversification from the microgeographic scale to a regional scale, we also included *O. algarvensis* individuals from three bays located around the island of Mallorca, which is approximately 800 km from Elba (see Figure 1b-d). We compared genomic divergence between Elba and Mallorca populations with local-level divergence around Elba, and we investigated the potential implications of microgeographic genomic differentiation in larger evolutionary time frames.

## Results and Discussion

This study aimed at understanding the potential for genomic diversification in a marine invertebrate that lacks dispersal capabilities. We used genomic data from hundreds of *O. algarvensis* individuals to assess population genetic divergence along the coastline of the island of Elba. Mitochondrial genomes of individuals from eight Elba bays showed a complex demographic history, while exomewide SNPs suggest that each bay harbors a reproductively isolated population. Geological forces that shaped habitat distributions during the Pleistocene are put forward as the main drivers of the population structure of *O. algarvensis* at a microgeographic scale today.

### The composition of mitochondrial COI gene haplogroups varies among locations at microgeographic and regional scales

To derive a mitochondrial phylogeography from the Elba and Mallorca locations, we screened the mitochondrial COI gene sequences of 1,406 individuals (Figure 2; Table 1a). We found a clear divergence between Mallorca and Elba-derived COI haplogroups, which was characterised by 31 nucleotide mutations within a 525 base pair alignment (Figure 2a; between haplogroups *h-g* and *a-e*). Within Elba, several haplogroups (*a-e*) are diverged by mutations between 1 and 5 nucleotides, but a divergent haplogroup *f* is separated from the rest of the Elba-derived haplogroups by 41 mutations (Figure 2a). The COI diversity in Elba was lowest in the northeast bays at Sorgente and Capo Vita that were dominated by the single haplogroup *a*, while Pomonte and Cavoli had a high COI diversity encompassing 3 to 5 dominant haplogroups, including the highly divergent haplogroup *f* (Figure 2b). This location-dependent phylogeographic pattern of COI haplogroups in Elba suggests that complex introgression events occurred in these populations. Potential mechanisms to form this pattern could include a regionwide spread of a haplogroup, site-dependent migration of novel haplogroup(s), or local extinction or survival in refugia of particular haplogroups during the glacial cycles seen in other fauna and flora (Médail and Diadema 2009; Dömel et al. 2015; Malekoutian et al. 2020). Using this distribution of the COI haplogroup diversity we selected sampling locations and the number of individuals for population genomic analyses. Genomic sequences of a total of 310 individuals from 11 locations were chosen for using low-coverage sequencing (Table 1b; average 1× coverage, average 9 million read pairs of 150 bp; sequences derived from the metagenome including the nuclear and mitochondrial genomes of *O. algarvensis* and its symbiotic bacteria (Sato et al. 2022)).

**Figure 2.**
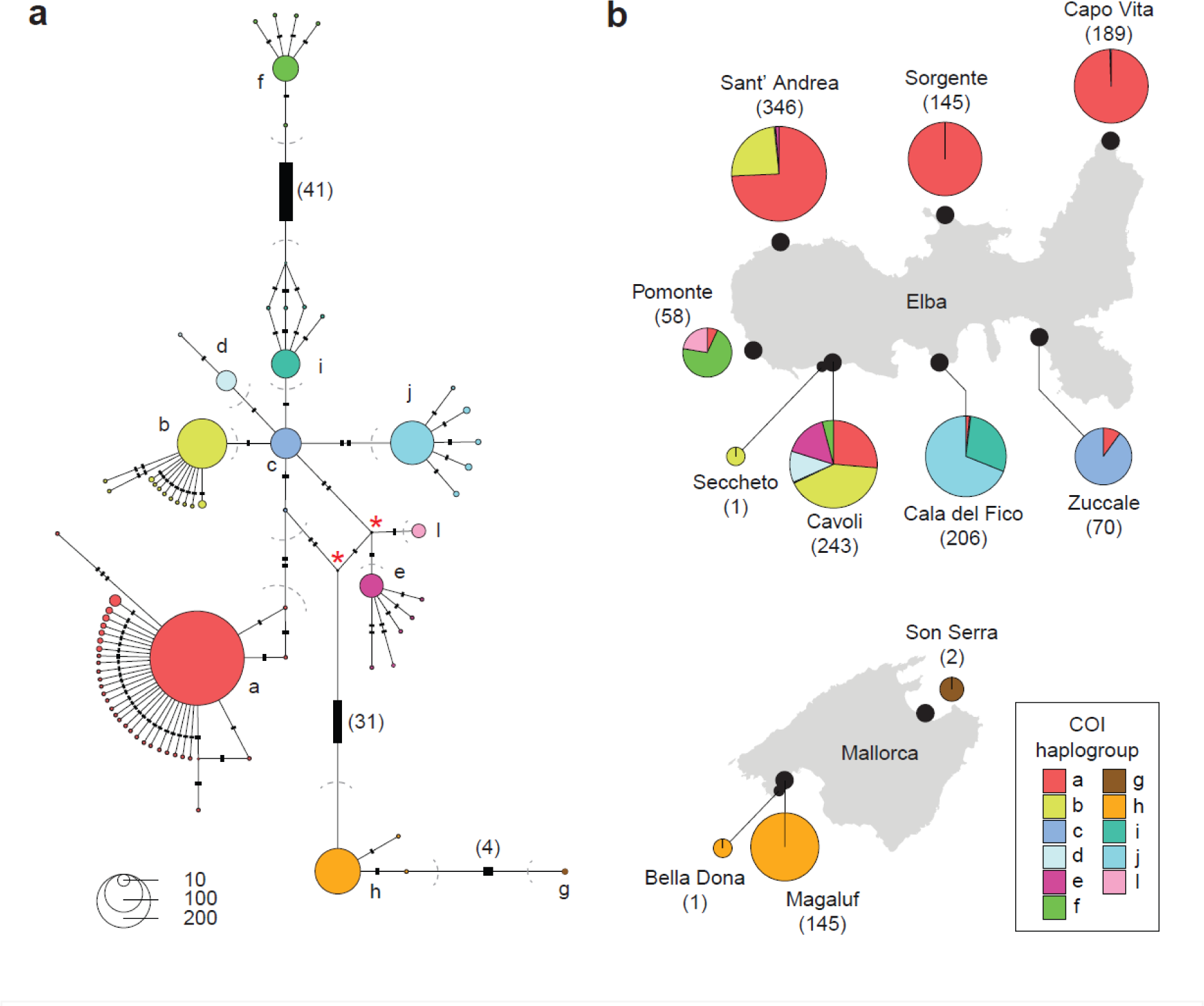
Distribution of the mitochondrial COI genes showed a clear mitochondrial divergence between Mallorca and Elba individuals and a complex introgression history in Elba. **(a)** Haplotype network of mitochondrial COI gene sequences based on 525 base pairs in a shared nucleotide alignment. Colours and alphabets indicate COI haplogroups, which are delimited by dashed lines. The size of circles corresponds to the number of samples identified. Hatch marks and associated numbers in brackets indicate the number of point mutations between haplotypes. Nodes indicated with a red asterisk indicate unobserved intermediates predicted by the haplotype network programme. (**b)** Distribution and relative abundances of haplogroups across the study sites. The size of pie-charts corresponds the total number of individuals studied per location shown in brackets.

**Table 1.**
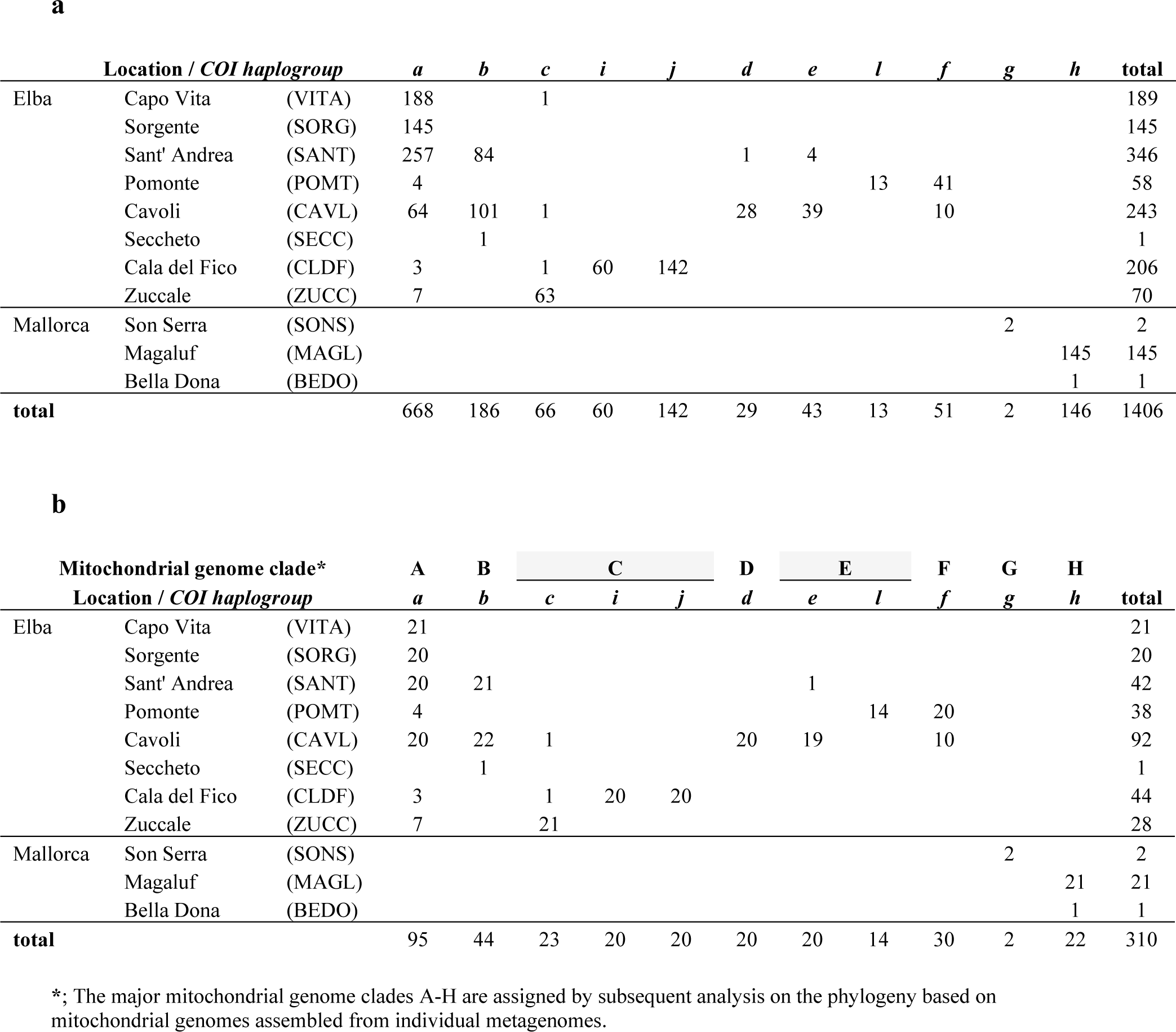
Overview of sample, locations, and mitochondrial lineages based on mitochondrial COI haplotypes and genomes. (**a**) Distribution of 1,406 individuals with mitochondrial COI haplogroups *a*-*h* based on 525 base pair amplicons across the study sites. (**b**) Number of replicated metagenomes generated from the samples above. The major mitochondrial genome clades A-H were assigned by subsequent phylogenetics on assembled, full mitochondrial genomes.

### Location-specific subclades of mitochondrial genomes within COI gene haplogroups suggest historical isolation between local populations

To further resolve mitochondrial phylogenies, we assembled complete circular mitochondrial genome sequences (mtDNA) from 219 individual metagenomes that contained sufficient mitochondrial read coverage. The phylogeny of the mtDNA showed three major divergent lineages corresponding to (i) the COI haplogroups *g* and *h* from Mallorca, (ii) COI haplogroup *f* from Elba, and (iii) all the rest of COI haplogroups from Elba (Figure 3). Within the third lineage, we identified five clades diverging by >0.4% average nucleotide differences, which we termed clades A – E (Figure 3b; Table 1b). Within the clades B, C, D and E, the mtDNA phylogeny showed monophyletic sister clades based on populations at Cavoli, Sant’ Andrea, Cala del Fico, Zuccale and Pomonte (Supplementary Figure S1). Monophyletic lineages within the clade A were also identified for the individuals from Capo Vita, Sorgente and Cala del Fico. This complete lineage sorting suggests that the migration of animals across bays on Elba has been restricted at the scale as small as 5 km. This also shows that the complex introgression indicated by multiple COI haplogroups per location was a historical event that occurred before such population isolation was established. In contrast, sub-clades within the clade A showed polyphyletic groups for individuals at each of Sant’ Andrea, Cavoli and Pomonte (Figure 3b, Supplementary Figure S1). These multiple sub-clades suggest that individuals within the mitochondrial clade A were widely distributed across the island and underwent multiple introgression and isolation events preceding the location-based separation. Furthermore, while a recent migration of individuals was likely restricted, incomplete lineage sorting in the clades A and F at Cavoli and Pomonte suggests that a rare migration was possible between these locations or from a shared unidentified origin at a contemporary time scale. It is worth noting that full mitochondrial sequences revealed location-specific subclades within some COI haplogroups (*a*, *b*, and *c*), which were not detectable in our COI marker gene survey. The retention of complex demographic history in mitochondrial marker sequences is well documented in smaller organisms with limited dispersal capabilities, such as marine meiofaunal taxa (Cerca et al. 2018), terrestrial annelids (Giska et al. 2015), estuarine fishes (Avise 1992; Jacobs et al. 2004), coastal echinoderms (Kelly and Palumbi 2010), molluscs (Kamel et al. 2014; Villamor et al. 2014), and kelp (Jacobs et al. 2004). The complex demographic history can then be attributed to population amalgamation, vicariance, and introgression resulting from tectonic movements, glacial cycles, and migration events (Avise 1992; Pelc et al. 2009; Kelly and Palumbi 2010). However, it is important to acknowledge that marker gene approaches often have limited temporal resolution due to the size of small sequence regions and their slow evolution rates. Our case study demonstrates that employing the complete mitochondrial genome sequence can further enhance our comprehension of mitochondrial patterns within populations, particularly by reconstructing demographic events at contemporary time scales.

**Figure 3.**
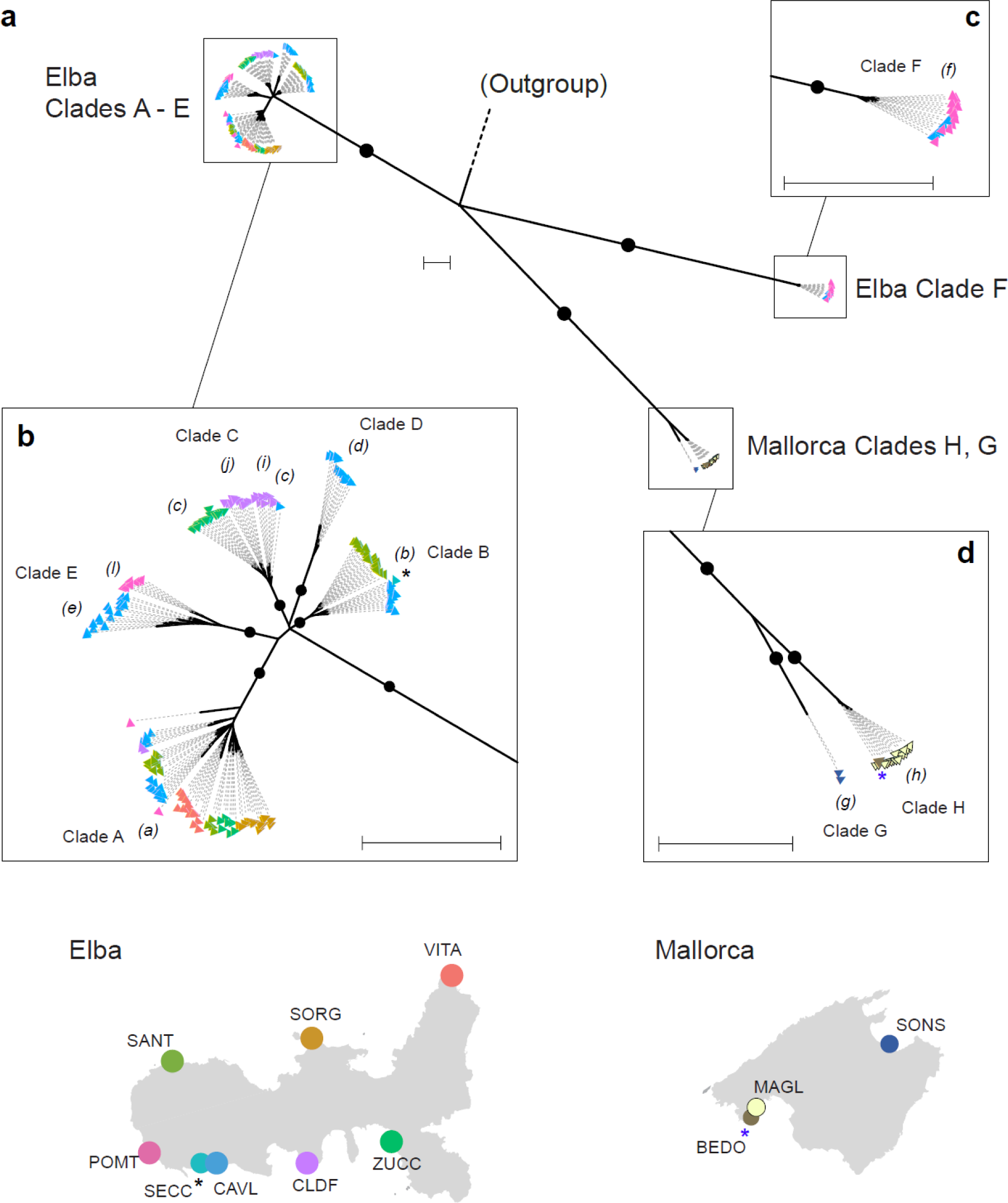
The phylogeny of mitochondrial genomes reveals major clades each showing location-based clustering within the islands. **(a)** Overall phylogeny of the mitochondrial genome clades. The colors of the tip labels correspond to their locations indicated on the maps at the bottom. Detailed phylogenies are indicated within insets for (**b**) Elba clades A-E, (**c**) Elba clade F, and (**d**) Mallorca clades G and H. Small italic alphabets in brackets indicate COI haplogroups. Posterior probability of clade support >0.95 are shown with black circles. For visibility, node supports within mitochondrial genome clades are provided in Supplementary Figure S1. All scales indicate 0.01 substitutions per base. Black and blue asterisks indicate Seccheto and Bella Donna, respectively, for ease of identifying their single occurrence each. Abbreviations for locations in the maps correspond to Table 1.

### Exomewide SNPs show distinct population structuring across locations at both microgeographic and regional scales

Mitochondrial data potentially indicate historical migration and allopatric divergence of individual mitochondrial lineages (Hellberg et al. 2002). However, they do not necessarily provide information about reproductive isolation or interbreeding between concurrent mitochondrial lineages (Ishida et al. 2011; Burton and Barreto 2012; Sloan et al. 2017). Investigations in this regard require data based on the nuclear genome. To examine population structuring, we investigated genetic divergence across *O. algarvensis* populations using exomewide SNPs. The genomic structuring across Elba populations indicated a clear clustering pattern corresponding to the locations that are as little as 5 km apart, with no overlap of genotype distributions (Figure 1c, Figure 4a, Supplementary Figure S2; PERMANOVA, p < 0.001; 376,768 SNPs). Only one exception was the Seccheto site as its genotype occurred within the cluster representing the population from Cavoli, 1 km away from Seccheto (Figure 4a). These results suggest that a coastal distance of 5 km can maintain genomic differentiation between populations of *O. algarvensis*, while a 1 km distance does not. Nonetheless, the fine spatial scale of isolation by 5 km despite the lack of a land-based physical barrier is remarkable. Population structuring at microgeographic scales (<10 km) have been reported in other marine invertebrates such as jellyfish (Dawson and Hamner 2005), mussels (de Leeuw et al. 2020), corals (Thomas et al. 2020), and sponges (Maas et al. 2023), but they are all associated with enclosed bodies of water like marine lakes, caves and lagoons. The passive dispersal of microscopic aquatic animals over long distances is plausible if these organisms have dormancy capabilities, resilience to the physical rigors of transportation, and the capacity for rapid colonization and reproduction (Fontaneto 2019). The incomplete lineage sorting of mitochondria in Cavoli and Pomonte populations suggests that long-distance dispersal indeed occurred. However, the microgeographic population structuring of nuclear genomes indicates that passive dispersal of *O. algarvensis* is so rare that it exerts negligible influence on the population’s genomic composition over ecological timescales. Such barriers to gene flow may similarly impact the population genetic structuring of numerous species with similar limited dispersal capacities. As argued by Dawson (2001), even a relatively short expanse of 100 kilometers of steep, rocky coastline can act as a significant phylogeographic barrier among coastal California marine fauna species with restricted dispersal abilities. The identification of gene flow barriers at small island scales may enable the detection of general patterns and their underlying geographical and environmental determinants.

**Figure 4.**
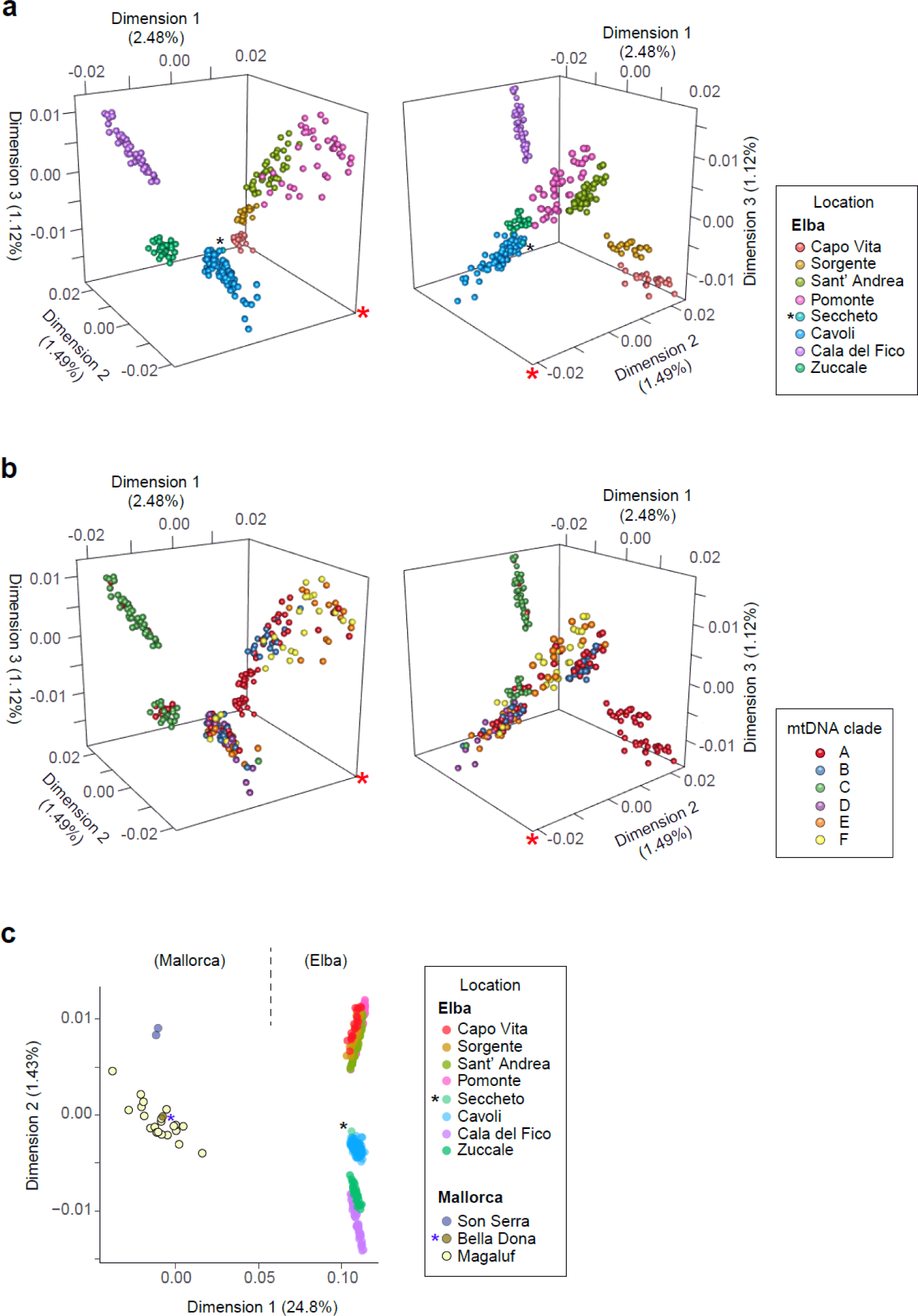
Population structuring based on exomewide SNPs showed clear genomic divergence explained by the locations between and within islands, but not by the mitochondrial lineages. MDSs based on genetic distances for individuals from Elba locations color-coded by locations (**a**) and the mitochondrial clades (**b**), and for individuals from all locations of Elba and Mallorca (**c**) and. The left and right 3D MDS plots in (**a**) and (**b**) are the identical plot that was horizontally rotated. The identical point of reference in the 3D space is indicated with red asterisks. Figures in brackets on the dimension axis labels indicate the proportion of variations explained by the dimension. Black and blue asterisks indicate Seccheto and Bella Donna, respectively, for ease of identifying their single occurrence each.

In contrast to the location-dependent population structure, nuclear genome population structure across Elba locations did not follow phylogenetic distances of the major mitochondrial genome clades (Figure 4b, Supplementary Figure S3). In other words, none of the major mitochondrial clades A – F was associated with a single cluster of the nuclear genome. Co-occurring mitochondrial lineages were not reflected as sub-clustering patterns of nuclear genomes within locations. This indicates that no reproductive isolation has been established between cooccurring mitochondrial lineages since their introduction and that their nuclear genomes have been well homogenised via genetic recombination. The divergent mitochondrial clade F, with 10% average nucleotide differences from the clades A-E, was not an exception, highlighting the discordance between the divergence of sympatric mitochondrial lineages and differentiation of nuclear genomes, like observed in e.g., snails (Thomaz et al. 1996), birds (Hogner et al. 2012), annelids (Giska et al. 2015) and orthopteran species (Morgan-Richards et al. 2017). Again, this underscores the necessity of nuclear genome analyses for understanding interbreeding population units. Inferring reproductively isolated populations based on high mitochondrial divergence only can be misleading.

In accordance with the population structuring across Elba locations at the microgeographic scale, we found a clear genomic divergence at a regional scale between populations of Mallorca and Elba separated by approximately 800 km (Figure 1b, Figure 4c; PERMANOVA, p < 0.001; 574,741 SNPs). The clustering of populations within the Mallorca region also illustrated that the northeast population (Son Serra) is genetically differentiated from the southeast populations (Magaluf and Bella Donna) that are approximately 150 km away in coastal distance (Figure 1d). Unlike the within-Elba comparison, the nuclear genome differentiation at these larger spatial scales was well aligned with the mitochondrial divergence (Figure 3a and 3d), indicating that population introgression was historically restricted between Mallorca and Elba, and between the northern and southern areas of Mallorca. On the other hand, no differentiation of nuclear genomes was present in individuals from Bella Donna and Magaluf that are 2 km apart (Figure 4c), suggesting that a barrier to gene flow can be maintained by a small distance between 2 km and 5 km (based on Mallorca and Elba populations, respectively).

### Isolation by distance explains population structuring in Elba, reflecting the coastline during the last glacial period

Comparisons of pairwise fixation indices (F_ST_) with geographic distances across Elba populations indicated that the population structure of *O. algarvensis* follows the pattern of isolation by distance (IBD; Figures 5a and 5b). This observation was supported by a Mantel test, which revealed a significant correlation (Mantel’s R = 0.514, p = 0.046) between pairwise F_ST_ values and distances.

**Figure 5.**
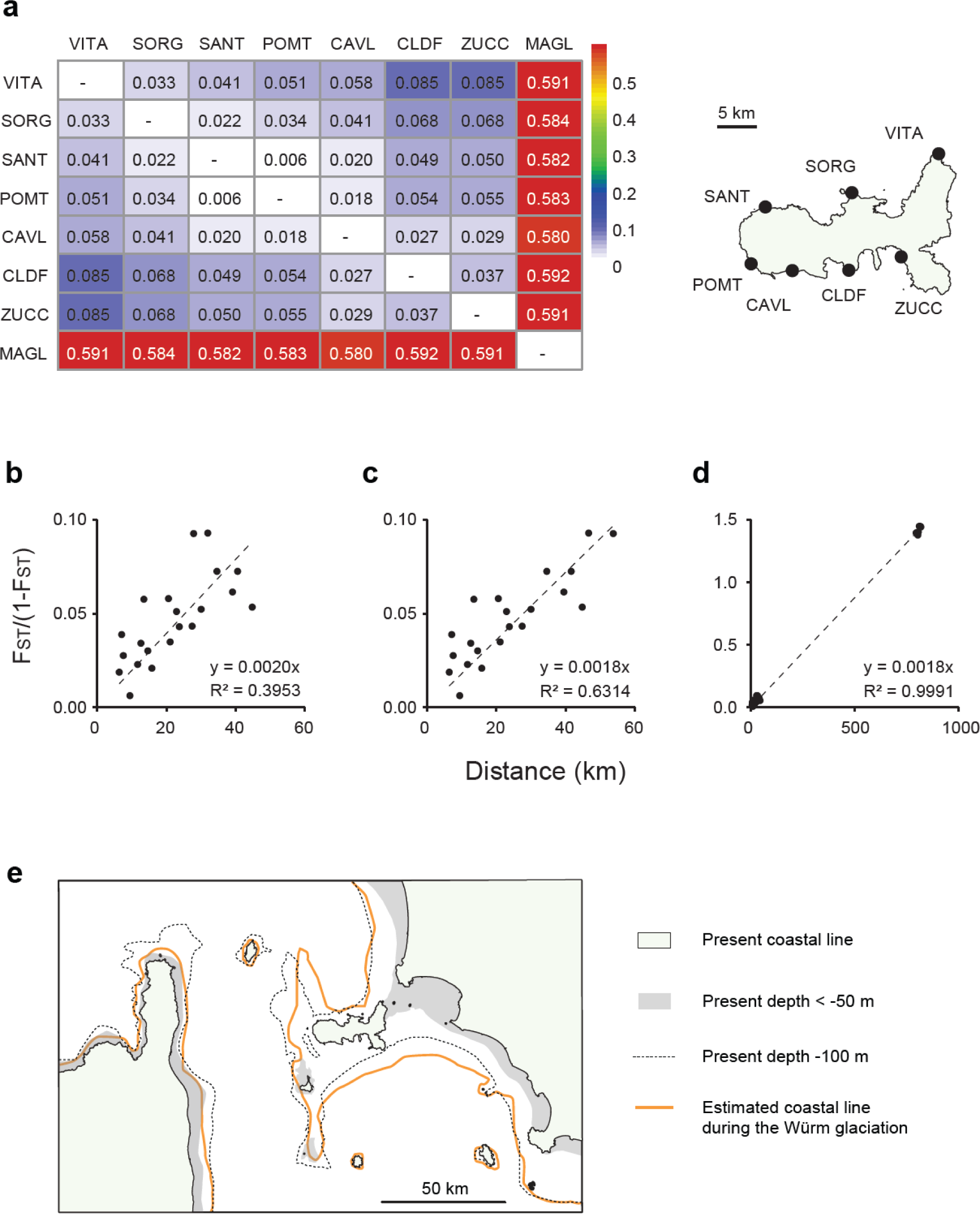
Pairwise FST and geographic distances across populations follow isolation by distance around Elba, reflecting the geological movement of coastlines during the last glacial cycles. (**a**) Heatmap of fixation indexes FST across study populations (Elba locations in the map, and Magaluf (MAGL) from Mallorca). (**b-d**) Scatter plots of pairwise geographic distances against linearised fixation index (FST/(1-FST)) for Elba populations (b), Elba populations considering a geographic barrier on the east coast of Elba (c), and all populations including ones from Elba and Mallorca (d), with the regression formula and squared correlation coefficient R2. (**e**) Coastal and benthic structures of the present and during the Würm glaciation in the Late Pleistocene (between ca. 115,000 and 11,700 years ago; according to the geological study by Bossio *et al*. (2000)).

However, there was one notable exception to this IBD pattern. The highest pairwise F_ST_ was observed between the north-eastern site of Capo Vita and two southern sites (Cala del Fico and Zuccale), despite these location pairs not having the greatest distances between locations (32 km and 28 km, respectively; cf. other pairs of Elba locations, such as Capo Vita and Pomonte with 45 km). Interestingly, a closer examination of topological data in the area revealed a shallow seafloor structure less than 50 m deep connecting the northeast of Elba to the Italian Peninsula (indicated by the grey area in Figure 5e). Geological evidence from sedimentary rock distribution suggests that this structure formed a land connection between the island and the mainland multiple times during the glacial cycles of the Pleistocene, when sea levels dropped more than 100 m below the present level (as depicted with the orange lines in Figure 5e; Bossio et al. 2000).

It is plausible to assume that this land connection acted as a barrier to gene flow between the northern and southern populations of *O. algarvensis* around Elba. When we accounted for this barrier in our analysis, we observed a remarkable improvement in fitting the IBD pattern (Figure 5c; Mantel’s R = 0.798, p = 0.006; Figure 5b). These findings underscore the significant influence of sedimental habitat positioning during the last glacial period on the current population structuring of *O. algarvensis* around Elba. Considering the link between seagrass meadows and *O. algarvensis* habitat (Giere and Erséus 2002), the fluctuations of sea level and corresponding shifts of seagrass habitats during the last glacial cycles are probable major drivers of the population structuring across Elba locations.

### The isolation by distance pattern extends to a regional scale between distant islands

The high values of pairwise F_ST_ revealed significant genetic divergence between the Mallorca populations and Elba populations, as depicted in Figure 4c and Figure 5a. To explore if the observed pattern of IBD at the microgeographic scale around Elba extends to explain the regional scale differences between these islands’ populations, we conducted further tests. Interestingly, we discovered that the slope of IBD, which represents the correlation coefficient between geographic distance and genetic distance, remained virtually identical when including both Elba and Mallorca populations (0.0018; Figures 5c and 5d). Bayesian models examining the co-regression between geographic distances and the F_ST_ at both regional and microgeographic scales revealed that the correlation coefficients were not significantly different from each other (difference = −0.00025 ∼ 0.00043, 95% highest density credibility intervals; Supplementary Text 1). These indistinguishable IBD slopes suggest a remarkably stable rate of population differentiation by distance across spatial scales, whereby the microgeographic IBD over a mere 60 km can account for the regional accumulation of genomic differentiation between populations separated by distances of up to 800 km (also see Supplementary Text 2). This finding underscores the substantial potential of retaining population genetic differentiation in *O. algarvensis*, as the microgeographic population divergence has consequences for genomic diversification over evolutionary timeframes.

Understanding the processes of speciation is a fundamental pursuit in unravelling the development of species richness within marine ecosystems (e.g., Palumbi 1994; Lessios et al. 2001; Butlin et al. 2008; Krug 2011; Roux et al. 2014). However, the initiation of speciation processes is challenging to study because of the time required for populations to accumulate genomic divergence and the fact that the ancestor shared by multiple species and the intermediates have become long extinct. Despite this challenge, we argue that new insights into the early driver of speciation may be obtained by examining within-species variations at microgeographic scales, and by exploring the stability of population differentiation across larger spatial scales. Our findings suggest that in non-dispersing marine interstitial organisms, genomic divergence can commence at surprisingly small spatial scales and accumulate consistently over substantial spatiotemporal dimensions. This phenomenon may be analogous to terrestrial organisms with limited dispersal capabilities, including flowering plants (Kisel and Barraclough 2010; Theim et al. 2014; Boucher et al. 2016) and large wingless insects (Dool et al. 2022).

### Conclusions

This study revealed population genetic divergence of a marine interstitial species with a high spatial resolution by leveraging an exomewide SNP analysis as a highly effective tool for detecting nuclear genome population structuring. Specifically, our results indicated that genetic structuring of non-dispersing invertebrates can occur at a microgeographic scale of less than 5 km. The use of complete circular mitochondrial genomes proved to be helpful in identifying barriers to gene flow across both historic and contemporary timescales. Along the dispersal limitation, local adaptation and founder effects have important roles in maintaining small-scale population differentiation require further studies (Orsini et al. 2013; Sexton et al. 2014; Wang and Bradburd 2014). However, dissecting local adaptation and genetic drifts is challenging in the presence of IBD. To investigate the potential selective pressures on a population, studies should involve characterizing the distribution of abiotic and biotic conditions as potential gradients, identify divergent loci along these gradients, and elucidate functional genomic regions with highly divergent alleles along the gradient.

Further work is desired to determine whether population genetic divergence across spatial scales, as observed in this study, is a common feature in other similar non-dispersal species because it may play a significant role in the genomic diversification of the whole community of marine interstitial invertebrates. Studying when and how fine-scale genetic divergence eventuates in speciation events in these organisms can provide further insights into the eco-evolutionary factors involved in the diversification of marine organisms.

## Materials and methods

### Study sites and sample collection

Specimens of *O. algarvensis* were collected around the islands of Elba, Italy and Mallorca, Spain, between 2013 and 2017. Samples were taken from eight bays around Elba and three bays around Mallorca (**Figure 1b-d**). *O. algarvensis* individuals were typically found in sandy sediment of mixed grain sizes (up to a few mm) in the vicinity of seagrass *Posidonia oceanica* at a water depth between 6 and 12 m. Sediments from the vertical layer between 5 and 30 cm deep were collected around the edge of seagrass meadows using a hand-held polyvinyl chloride core (custom made; 20 cm in diameter, 30 cm long). Collected sediments were carefully contained into a lidded 15L bucket underwater, and the bucket was immediately transported to an indoor facility. *O. algarvensis* individuals were sorted from the sediment using a series of seawater rinses and filtering onto 150 µm meshes. Sorted specimens were individually fixed in 500 µl of RNAlater (Sigma-Aldrich, Darmstadt, Germany) and stored at 4°C until DNA extraction.

### Extraction of DNA and screening of mitochondrial genes

DNA was extracted from a total of 1,406 *O. algarvensis* individuals using the DNeasy 96 Blood & Tissue Kit (Qiagen; Hilden, Germany) according to the manufacturer’s instructions. To verify the morphological species identification of the individual specimens, the mitochondrial cytochrome c oxidase subunit I (COI) gene was PCR-amplified using the primer set of COI-1490F and COI-2198R and Sanger-sequenced as previously described (Folmer et al. 1994). Obtained COI sequences were aligned with reference COI-sequences of gutless oligochaetes using MAFFT v1.5.0 multiple-alignment algorithm (Katoh and Standley 2013), and their phylogenetic affiliation to *O. algarvensis* was verified with a phylogenetic tree drawn with IQ-TREE v2.1.2 using default parameters (Nguyen et al. 2015). Species identification was also confirmed with the identical 18S rRNA gene sequences extracted from metagenomes (see below). A COI haplotype network was built based on a 525bp long, trimmed sequence alignment of all *O. algarvensis* individuals, using PopART v1.7 (Leigh and Bryant 2015) with the statistical parsimony method (TCS; Clement et al. 2000).

### DNA library preparation and sequencing

We obtained metagenomes of a total of 310 *O. algarvensis* individuals, representing different combinations of locations and mitochondrial COI-lineages (**Table 1b**). Sequencing libraries of the individual DNA samples were generated by following a cost-effective, multiplexed library preparation method using a Tn5 transposase purification and tagmentation protocol (Hennig et al. 2018). The Tn5 transposase was produced by the Protein Science Facility at Karolinska Institutet SciLifeLab (http://ki.se/psf), Stockholm, Sweden. Quantity and quality of DNA samples were checked with (i) the Quantus Fluorometer with the QuantiFluor dsDNA System, (ii) Agilent TapeStation System with DNA ScreenTape, and (iii) FEMTO Pulse genomic DNA analysis (Promega Corporation, Madison, WI, USA; Agilent Technologies, Santa Clara, CA, USA; Advanced Analytical Technologies Inc., Heidelberg, Germany, respectively). Insert template DNA was size-selected targeting 400 - 500 bp using AMPure XP (Beckman Coulter, Indianapolis, IN, USA), and paired-end sequences of 150 bp x 2 were generated using the Illumina HiSeq3000 System (Illumina, San Diego, CA, USA), targeting a yield of 2 gigabases data per sample. Library-preparation, quality control, and sequencing were performed at the Max Planck-Genome-Centre Cologne (MP-GC, Cologne, Germany).

### Transcriptomic reference assembly

A transcriptome reference of *O. algarvensis* was assembled from total RNA sequences derived from 12 individual animals from Sant’ Andrea Bay in Elba, which were maintained under experimental anoxic and oxic conditions to increase the diversity of transcripts. Six animals were incubated for 24 hours in anoxic artificial seawater in gas-tight serum bottles and fixed in RNAlater (‘anoxic worms’), while another six animals were also incubated first in anoxic artificial seawater for 24 hours in the same manner but subsequently kept in fully oxygenated artificial seawater for 12 hours before fixation (‘oxic worms’). Total RNA was extracted with the AllPrep DNA/RNA kit (Qiagen), following the default protocol with a 3 min bead-beating step at 20 Hz. Quantity and quality of RNA samples were checked with the Bioanalyzer 2100 (Agilent Technologies). Synthesis of cDNA was performed with the Ovation RNA-Seq System (NuGEN Technologies Inc., Redwood City, CA, USA), and DNA shearing was applied with the Covaris ultrasonicator (Covaris, Woburn, MA, USA). Sequencing libraries were prepared with the NEBNext Ultra DNA Library Prep Kit for Illumina (New England Biolabs, Ipswich, MA, USA), targeting the insert size of 250 bp. Metatranscriptomic reads of approximately 6 gigabases (paired-end reads of 100 bases x 2; 30 M pairs) were retrieved from each of 12 *O. algarvensis* animals, using the HiSeq2500 System (Illumina) at the MP-GC.

A high quality transcriptomic reference of *O. algarvensis* was assembled *de novo* with a workflow built upon on the guideline by De Wit *et al*. (2015) but modified to accommodate input metatranscriptomic sequences from a symbiotic organism (Supplementary Figure S4 ‘transcript reference assembly workflow’). Briefly, the goal of the workflow is to generate a set of transcriptomes, each of which corresponds to a single genomic region of the host (i.e., isoforms and allelic variations are clustered and have a single representative sequence each, but paralogs are separated, and do not contain transcripts from other organisms such as endosymbionts). Each of the metatranscriptomic datasets was first de-contaminated from residual Illumina adapters and PhiX sequences and quality filtered using bbduk (BBTools v37.86; https://jgi.doe.gov/data-and-tools/bbtools/). Sequencing errors were corrected using SEECER v0.1.3 (Le et al. 2013), and sequences of *O. algarvensis* mitochondrial genome DNA (mtDNA) and symbiotic bacteria, ribosomal RNA (rRNA), and known DNA contaminants (e.g. *Drosophila* sp.) were removed using bbmap (BBTools). For mapping references, we used mtDNA and symbiont genome bins previously assembled from *O. algarvensis* (Woyke et al. 2006; Sato et al. 2022), all rRNA sequences from sortMeRNA databases v4.3.4 (Kopylova et al. 2012), and the *Drosophila melanogaster* genome (GenBank assembly accession: GCA_000001215.4). Clean reads were pooled and co-assembled *de novo* using Trinity v2.4.0 (Grabherr et al. 2011). Contigs of < 600 bases, < 4× coverage, or without open reading frames (ORFs) identified with TransDecoder v3.0.1 (Haas et al. 2013) were filtered out to remove most of the assembly artifacts. To remove contaminating viral, bacterial, archaeal, and other eukaryotic (e.g., Cnidaria, Viridiplantae) sequences, taxonomic affiliations of contigs were identified using DIAMOND blastx search v0.8.36 (Buchfink et al. 2014) against the NCBI-nr protein database (https://www.ncbi.nlm.nih.gov/), and only contigs identified within the clade Bilateria (encompassing the phylum Annelida) were exported using MEGAN v6.19.6 (Huson et al. 2007). Redundant contigs due to structurally similar isoforms and allelic variations among *O. algarvensis* individuals were removed with Corset v1.06 (Davidson and Oshlack 2014), with the reads from the ‘oxic’ and ‘anoxic worms’ being specified as separate treatment groups. Finally, differentially spliced isoforms were clustered using Trinity isoform tags by choosing the longest contigs per gene. Completeness of the transcriptomic assembly was assessed at each filtering stage with BUSCO v2.0.1 (Simão et al. 2015) using the metazoan single copy orthologous gene database (odb9) built in BUSCO v2.0.1.

### Identification of exomewide SNPs

Exomewide SNPs were identified among *O. algarvensis* individuals based on low-coverage genomewide sequencing using an *in silico* exome capturing workflow (Therkildsen and Palumbi 2016). We made a few modifications for (i) processing metagenomic data recovered from the host animal and abundant endosymbionts and (ii) increasing the efficiency of avoiding false-positive SNP-sites derived from repeat regions in the exome. Briefly, overall genomic contents of raw reads were first checked by assembling and cataloguing small subunit ribosomal RNA genes (16S rRNA and 18S rRNA) using PhyloFlash v3.3b (Gruber-Vodicka et al. 2020). After removing exact PCR-derived sequence duplications and residual Illumina adapter barcode sequences, genomic reads matching the *O. algarvensis* symbionts and mitochondria sequences were removed using reference genomes that were publicly available (Woyke et al. 2006; Sato et al. 2022) and newly assembled in this study using the method previously described (Sato et al. 2022). After read quality-filtering, merging of overlapping paired-end reads, removal of ribosomal and human-, viral-, and bacterial-contaminant sequences, the remaining data was assumed to contain only *O. algarvensis* genomic sequences. Input data were size-capped to 1.8 gigabases per sample with the minimum of 1.0 gigabases to ensure similar mapping depths among samples and improve the accuracy of SNP-detection and downstream analyses. The final clean reads were mapped to the transcriptomic reference, followed by the mapping-quality based filtering (Therkildsen and Palumbi 2016), which resulted in an average mapping depth of approximately 1.0× across the whole exome. SNPs in the exome were identified by computing posterior genotype probabilities in a Bayesian approach with ANGSD v0.929 (Nielsen et al. 2012; Korneliussen et al. 2014), using Hardy Weinberg equilibrium applied to sample allele frequency (SAF) as a prior.

### Population genetic structures

To exclude erroneous identification of SNPs derived from genetic repeat regions (i.e., a potential source of genotyping errors), the maximum individual coverage depth of 4 was specified in ANGSD (-setMaxIndDepth) after examining the frequency distribution of read depths. SNP-sites were also filtered with a mapping depth range covered by between 5 and 80 percentiles of reads (-setMinDepth, -setMaxDepth) and covered in >50% of individuals analysed (-minInd). To further increase the reliability of SNP identification, we generated a region filter file (-r) based on the contig coverage data of random 30 samples. This filter contained contigs that the mean read coverage was less than 10× but excluded contigs that had no reads mapped or all of samples indicated >4× coverages (this setting effectively excluded contigs that attracted reads from repeat regions or erroneously clustered paralogues but allowed high coverage sites in some samples due to stochasticity). SNP-sites using a *p*-value cutoff of 0.001 (-SNP_pval) and minimum minor allele frequency of 0.01 (-minMaf) were retained for population structure analyses. Based on the genotype probabilities at these SNP-sites, pairwise genetic distances between individual animals were obtained with ngsDist v1.0.2 (Vieira et al. 2016) and visualised on multi-dimensional scaling (MDS) plots using *ggplot2* v3.4.0 (Wickham 2016) and *rgl* v0.110.2 (Murdoch et al. 2022) packages implemented in the statistical environment R v3.6.2 (R_Core_Team 2016). Clustering patterns in MDS in relation to the sample locations and mitochondrial lineages were examined using permutational multivariate analysis of variance (PERMANOVA) and pairwise PERMANOVA, implemented in R-packages *vegan* v2.6 (Oksanen et al. 2017) and *RVAideMemoire* v0.9-81.2 (Hervé 2022), respectively.

### Population genetic summary statistics

The population genetic differentiation metric (fixation index; F_ST_) was computed using 20 random individual animals per location. Seccheto, Bella Donna and Son Serra samples were excluded from the F_ST_ analysis as the sample numbers were insufficient (n = 1 or 2 individuals per location). Pairwise global F_ST_ between populations were estimated from the site frequency spectrum (SFS) based on the SAF likelihoods instead of genotype calling using ANGSD programs *realSFS* (Nielsen et al. 2012; Korneliussen et al. 2013). Unfolded SFS estimates were used for F_ST_ calculations by polarising alleles with the transcriptome reference and constructing pairwise 2D-SFSs.

To assess the spatial scale of population genetic differentiation, isolation by distance (IBD) analyses were performed on the regression of population genetic differentiation (linearised F_ST;_ F_ST_ / (1-F_ST_)) against sea distance between population origins using Mantel tests (Mantel 1967; Rousset 1997). Mantel tests were performed with 999 permutations, using the *mantel* function in the R-package *vegan* (Oksanen et al. 2017). Sea distances were acquired by a path connecting closest sea distances between locations using the ruler-path tool in Google Earth Pro v7.3 (https://www.google.com/earth). The bathymetric contour at -100 m indicated in Figure 5 was based in the Navionics Chart Viewer (available at https://webapp.navionics.com).

### Mitochondrial genome genealogy

Circular full mitochondrial genome assemblies were obtained using the method previously described (Sato et al. 2022), and the directions of circular sequences were checked in Geneious 11.0.3 (https://www.geneious.com). Unidirectional sequences were aligned using MAFFT (Katoh and Standley 2013), and phylogenetic relationships of mtDNA sequences were constructed from the alignment using the MR-Bayes v2.2.4 (Ronquist et al. 2012) and visualised in iTOL v4.0 (Letunic and Bork 2019).

## Data Availability

Sequence data generated in this study are available in the European Nucleotide Archive (ENA) under accession number PRJEB55913. Annotated bioinformatic scripts and reference files are available in a GitHub depository (https://github.com/yuisato/Oalg_exome_snps).

## Acknowledgements

We thank Miriam Weber, Christian Lott, staff of the HYDRA Institute, Ramon Rosselló-Móra, the Institut Mediterranean d’Estudis Avançats, Francois Seneca of the Scientific Centre of Monaco, Jean Seneca, Patricia Warner, Katrin Schmidt, and members of the Symbiosis Department at the Max Planck Institute for Marine Microbiology for their assistance in sample collection. We also thank Iliana Baums (Penn State University) for insightful discussion, Thorfinn Korneliussen (University of Copenhagen) and Murray Logan (Australian Institute of Marine Science) for technical advice on analyses. This study was supported by a Gordon and Betty Moore Foundation Marine Microbial Initiative Investigator Award (grant number GBMF3811) and the Max Planck Society. The authors declare no conflict of interest.

## References

Andrews KR, Good JM, Miller MR, Luikart G, Hohenlohe PA. 2016. Harnessing the power of RADseq for ecological and evolutionary genomics. Nature Reviews Genetics 17:81–92.

Arroyo NL, Aarnio K, Bonsdorff E. 2006. Drifting algae as a means of re-colonizing defaunated sediments in the Baltic Sea. A short-term microcosm study. Hydrobiologia 554:83–95.

Avise JC. 1992. Molecular population structure and the biogeographic history of a regional fauna: A case history with lessons for conservation biology. Oikos 63:62–76.

Bossio A, Cornamusini G, Ferrandini J, Ferrandini M, Maria Foresi L, Mazzanti R, Mazzei R, Salvatorini G, Sandrelli F. 2000. Dinamica dal Neogene al Quaternario della Corsica orientale e della Toscana. In. L’attività scientifica delle Università di Pisa e Corte. Pisa: Edizioni ETS. p. 87–95.

Boucher FC, Zimmermann NE, Conti E. 2016. Allopatric speciation with little niche divergence is common among alpine Primulaceae. Journal of Biogeography 43:591–602.

Bradbury IR, Laurel BJ, Snelgrove PVR, Bentzen P, Campana SE. 2008. Global patterns in marine dispersal estimates: the influence of geography, taxonomic category and life history. Proceedings of the Royal Society B: Biological Sciences 275:1803–1809.

Buchfink B, Xie C, Huson DH. 2014. Fast and sensitive protein alignment using DIAMOND. Nature Methods 12:59.

Burton RS, Barreto FS. 2012. A disproportionate role for mtDNA in Dobzhansky–Muller incompatibilities? Molecular Ecology 21:4942–4957.

Butlin RK, Galindo J, Grahame JW. 2008. Sympatric, parapatric or allopatric: the most important way to classify speciation? Philosophical Transactions of the Royal Society B: Biological Sciences 363:2997–3007.

Cerca J, Purschke G, Struck TH. 2018. Marine connectivity dynamics: clarifying cosmopolitan distributions of marine interstitial invertebrates and the meiofauna paradox. Marine Biology 165:123.

Chandler GT, Fleeger JW. 1983. Meiofaunal colonization of azoic estuarine sediment in Louisiana: Mechanisms of dispersal. Journal of Experimental Marine Biology and Ecology 69:175–188.

Clement M, Posada D, Crandall KA. 2000. TCS: a computer program to estimate gene genealogies. Molecular Ecology 9:1657–1659.

Curini-Galletti M, Artois T, Delogu V, De Smet WH, Fontaneto D, Jondelius U, Leasi F, Martínez A, Meyer-Wachsmuth I, Nilsson KS, et al. 2012. Patterns of Diversity in Soft-Bodied Meiofauna: Dispersal Ability and Body Size Matter. PLOS ONE 7:e33801.

Davidson NM, Oshlack A. 2014. Corset: enabling differential gene expression analysis for de novo assembled transcriptomes. Genome Biology 15:410.

Dawson MN. 2001. Phylogeography in coastal marine animals: a solution from California? Journal of Biogeography 28:723–736.

Dawson MN, Hamner WM. 2005. Rapid evolutionary radiation of marine zooplankton in peripheral environments. Proceedings of the National Academy of Sciences 102:9235–9240.

de Leeuw CA, Peijnenburg KTCA, Gillespie RG, Maas DL, Hanzawa N, Tuti Y, Toha AHA, Aji LP, Becking LE. 2020. First come, first served: Possible role for priority effects in marine populations under different degrees of dispersal potential. Journal of Biogeography 47:1649–1662.

De Wit P, Pespeni MH, Palumbi SR. 2015. SNP genotyping and population genomics from expressed sequences – current advances and future possibilities. Molecular Ecology 24:2310–2323.

Derycke S, Remerie T, Backeljau T, Vierstraete A, Vanfleteren J, Vincx M, Moens T. 2008. Phylogeography of the *Rhabditis* (*Pellioditis*) *marina* species complex: evidence for long-distance dispersal, and for range expansions and restricted gene flow in the northeast Atlantic. Molecular Ecology 17:3306–3322.

Dömel JS, Convey P, Leese F. 2015. Genetic data support independent glacial refugia and open ocean barriers to dispersal for the Southern Ocean Sea Spider *Austropallene cornigera* (Möbius, 1902). Journal of Crustacean Biology 35:480–490.

Dool SE, Picker MD, Eberhard MJB. 2022. Limited dispersal and local adaptation promote allopatric speciation in a biodiversity hotspot. Molecular Ecology 31:279–295.

Folmer O, Black M, Hoeh W, Lutz R, Vrijenhoek R. 1994. DNA primers for amplification of mitochondrial cytochrome c oxidase subunit I from diverse metazoan invertebrates. Molecular marine biology and biotechnology 3:294–299.

Fontaneto D. 2019. Long-distance passive dispersal in microscopic aquatic animals. Movement Ecology 7:10.

Fuentes-Pardo AP, Ruzzante DE. 2017. Whole-genome sequencing approaches for conservation biology: Advantages, limitations and practical recommendations. Molecular Ecology 26:5369–5406.

Giere O. 2006. Ecology and biology of marine Oligochaeta – an inventory rather than another review. Hydrobiologia 564:103–116.

Giere O. 2009. Meiobenthology: The microscopic motile fauna of aquatic sediments. Berlin Heidelberg: Springer-Verlag.

Giere O, Erséus C. 2002. Taxonomy and new bacterial symbioses of gutless marine Tubificidae (Annelida, Oligochaeta) from the Island of Elba (Italy). Organisms Diversity & Evolution 2:289–297.

Giere O, Erséus C, Stuhlmacher F. 1998. A new species of Olavius (Tubificidae) from the Algarve Coast in Portugal, the first East Atlantic gutless oligochaete with symbiotic bacteria. Zoologischer Anzeiger 237:209–214.

Giska I, Sechi P, Babik W. 2015. Deeply divergent sympatric mitochondrial lineages of the earthworm *Lumbricus rubellus* are not reproductively isolated. BMC Evolutionary Biology 15:217.

Grabherr MG, Haas BJ, Yassour M, Levin JZ, Thompson DA, Amit I, Adiconis X, Fan L, Raychowdhury R, Zeng Q, et al. 2011. Full-length transcriptome assembly from RNA-Seq data without a reference genome. Nature Biotechnology 29:644.

Gruber-Vodicka HR, Seah BKB, Pruesse E. 2020. phyloFlash: Rapid small-subunit rRNA profiling and targeted assembly from metagenomes. mSystems 5:e00920–00920.

Haas BJ, Papanicolaou A, Yassour M, Grabherr M, Blood PD, Bowden J, Couger MB, Eccles D, Li B, Lieber M, et al. 2013. De novo transcript sequence reconstruction from RNA-seq using the Trinity platform for reference generation and analysis. Nature protocols 8:1494–1512.

Hagerman GM, Rieger RM. 1981. Dispersal of benthic meiofauna by wave and current action in Bogue Sound, North Carolina, USA. Marine Ecology 2:245–270.

Hellberg ME, Burton RS, Neigel JE, Palumbi SR. 2002. Genetic assessment of connectivity among marine populations. Bulletin of Marine Science 70:273–290.

Hennig BP, Velten L, Racke I, Tu CS, Thoms M, Rybin V, Besir H, Remans K, Steinmetz LM. 2018. Large-scale low-cost NGS library preparation using a robust Tn5 purification and tagmentation protocol. G3 (Bethesda) 8:79–89.

Hervé M. 2022. RVAideMemoire: Testing and plotting procedures for biostatistics. https://cran.r-project.org/package=RVAideMemoire.

Hogner S, Laskemoen T, Lifjeld JT, Porkert J, Kleven O, Albayrak T, Kabasakal B, Johnsen A. 2012. Deep sympatric mitochondrial divergence without reproductive isolation in the common redstart Phoenicurus phoenicurus. Ecology and Evolution 2:2974–2988.

Huson DH, Auch AF, Qi J, Schuster SC. 2007. MEGAN analysis of metagenomic data. Genome Research 17:377–386.

Ishida Y, Oleksyk TK, Georgiadis NJ, David VA, Zhao K, Stephens RM, Kolokotronis S-O, Roca AL. 2011. Reconciling apparent conflicts between mitochondrial and nuclear phylogenies in African elephants. PLOS ONE 6:e20642.

Jacobs DK, Haney TA, Louie KD. 2004. Genes, diversity, and geologic process on the Pacific coast. Annual Review of Earth and Planetary Sciences 32:601–652.

Kamel SJ, Grosberg RK, Addison JA. 2014. Multiscale patterns of genetic structure in a marine snail (*Solenosteira macrospira*) without pelagic dispersal. Marine Biology 161:1603–1614.

Katoh K, Standley DM. 2013. MAFFT multiple sequence alignment software version 7: Improvements in performance and usability. Molecular Biology and Evolution 30:772–780.

Kelly RP, Palumbi SR. 2010. Genetic structure among 50 species of the Northeastern Pacific rocky intertidal community. PLOS ONE 5:e8594.

Kieneke A, Nikoukar H. 2017. Integrative morphological and molecular investigation of Turbanella hyalina Schultze, 1853 (Gastrotricha: Macrodasyida), including a redescription of the species. Zoologischer Anzeiger 267:168–186.

Kisel Y, Barraclough TG. 2010. Speciation has a spatial scale that depends on levels of gene flow. The American Naturalist 175:316–334.

Kopylova E, Noé L, Touzet H. 2012. SortMeRNA: fast and accurate filtering of ribosomal RNAs in metatranscriptomic data. Bioinformatics 28:3211–3217.

Korneliussen TS, Albrechtsen A, Nielsen R. 2014. ANGSD: Analysis of next generation sequencing data. BMC Bioinformatics 15:356.

Korneliussen TS, Moltke I, Albrechtsen A, Nielsen R. 2013. Calculation of Tajima’s D and other neutrality test statistics from low depth next-generation sequencing data. BMC Bioinformatics 14:289.

Krug PJ. 2011. Patterns of speciation in marine gastropods: A review of the phylogenetic evidence for localized radiations in the sea. American Malacological Bulletin 29:169–186, 118.

Le H-S, Schulz MH, McCauley BM, Hinman VF, Bar-Joseph Z. 2013. Probabilistic error correction for RNA sequencing. Nucleic Acids Research.

Leigh JW, Bryant D. 2015. popart: full-feature software for haplotype network construction. Methods in Ecology and Evolution 6:1110–1116.

Lessios HA, Kessing BD, Pearse JS. 2001. Population structure and speciation in tropical seas: Global phylogeography of the sea urchin Diadema. Evolution 55:955–975.

Letunic I, Bork P. 2019. Interactive Tree Of Life (iTOL) v4: recent updates and new developments. Nucleic Acids Research 47:W256–W259.

Maas DL, Prost S, de Leeuw CA, Bi K, Smith LL, Purwanto P, Aji LP, Tapilatu RF, Gillespie RG, Becking LE. 2023. Sponge diversification in marine lakes: Implications for phylogeography and population genomic studies on sponges. Ecology and Evolution 13:e9945.

Mackiewicz P, Matosiuk M, Świsłocka M, Zachos FE, Hajji GM, Saveljev AP, Seryodkin IV, Farahvash T, Rezaei HR, Torshizi RV, et al. 2022. Phylogeny and evolution of the genus *Cervus* (Cervidae, Mammalia) as revealed by complete mitochondrial genomes. Scientific Reports 12:16381.

Malekoutian M, Sharifi M, Vaissi S. 2020. Mitochondrial DNA sequence analysis reveals multiple Pleistocene glacial refugia for the Yellow-spotted mountain newt, Neurergus derjugini (Caudata: Salamandridae) in the mid-Zagros range in Iran and Iraq. Ecology and Evolution 10:2661–2676.

Mantel N. 1967. The Detection of Disease Clustering and a Generalized Regression Approach. Cancer Research 27:209–220.

Médail F, Diadema K. 2009. Glacial refugia influence plant diversity patterns in the Mediterranean Basin. Journal of Biogeography 36:1333–1345.

Morgan-Richards M, Bulgarella M, Sivyer L, Dowle EJ, Hale M, McKean NE, Trewick SA. 2017. Explaining large mitochondrial sequence differences within a population sample. Royal Society Open Science 4:170730.

Murdoch D, Adler D, Nenadic O, Urbanek S, Chen M, Gebhardt A, Bolker B, Csardi G, Strzelecki A, Senger A, et al. 2022. Package ‘rgl’: 3D Visualization Using OpenGL. https://github.com/dmurdoch/rgl.

Narjes S, Jelle VC, Annelien R, Hadi M, Frederik L, Tom M. 2017. Lack of population genetic structure in the marine nematodes Ptycholaimellus pandispiculatus and Terschellingia longicaudata in beaches of the Persian Gulf, Iran. Marine Ecology 38:e12426.

Nguyen L-T, Schmidt HA, von Haeseler A, Minh BQ. 2015. IQ-TREE: A Fast and Effective Stochastic Algorithm for Estimating Maximum-Likelihood Phylogenies. Molecular Biology and Evolution 32:268–274.

Nielsen R, Korneliussen T, Albrechtsen A, Li Y, Wang J. 2012. SNP calling, genotype calling, and sample allele frequency estimation from new-generation sequencing data. PLOS ONE 7:e37558.

Oksanen J, Blanchet FG, Friendly M, Kindt R, Legendre P, McGlinn D, Minchin PR, O’Hara RB, Simpson GL, Solymos P, et al. 2017. vegan: Community ecology package. R package version 2.4–4. https://CRAN.R-project.org/package=vegan.

Orsini L, Vanoverbeke J, Swillen I, Mergeay J, De Meester L. 2013. Drivers of population genetic differentiation in the wild: isolation by dispersal limitation, isolation by adaptation and isolation by colonization. Molecular Ecology 22:5983–5999.

Palmer MA. 1986. Hydrodynamics and structure: Interactive effects on meiofauna dispersal. Journal of Experimental Marine Biology and Ecology 104:53–68.

Palumbi SR. 1994. Genetic divergence, reproductive isolation, and marine speciation. Annual Review of Ecology and Systematics 25:547–572.

Pelc RA, Warner RR, Gaines SD. 2009. Geographical patterns of genetic structure in marine species with contrasting life histories. Journal of Biogeography 36:1881–1890.

Presgraves DC. 2010. The molecular evolutionary basis of species formation. Nature Reviews Genetics 11:175–180.

R_Core_Team. 2016. R: A language and environment for statistical computing. Vienna, Austria: R Foundation for Statistical Computing.

Radziejewska T, Gruszka P, Rokicka-Praxmajer J. 2006. A home away from home: A meiobenthic assemblage in a ship’s ballast water tank sediment. Oceanologia 48:259–265.

Ronquist F, Teslenko M, van der Mark P, Ayres DL, Darling A, Höhna S, Larget B, Liu L, Suchard MA, Huelsenbeck JP. 2012. MrBayes 3.2: efficient Bayesian phylogenetic inference and model choice across a large model space. Systematic Biology 61:539–542.

Rousset F. 1997. Genetic Differentiation and Estimation of Gene Flow from F-Statistics Under Isolation by Distance. Genetics 145:1219–1228.

Roux C, Fraïsse C, Castric V, Vekemans X, Pogson GH, Bierne N. 2014. Can we continue to neglect genomic variation in introgression rates when inferring the history of speciation? A case study in a Mytilus hybrid zone. Journal of Evolutionary Biology 27:1662–1675.

Sato Y, Wippler J, Wentrup C, Ansorge R, Sadowski M, Gruber-Vodicka H, Dubilier N, Kleiner M. 2022. Fidelity varies in the symbiosis between a gutless marine worm and its microbial consortium. Microbiome 10:178.

Savolainen O, Lascoux M, Merila J. 2013. Ecological genomics of local adaptation. Nature Reviews Genetics 14:807–820.

Schmidt H, Westheide W. 2000. Are the meiofaunal polychaetes *Hesionides arenaria* and *Stygocapitella subterranea* true cosmopolitan species? — results of RAPD-PCR investigations. Zoologica Scripta 29:17–27.

Schmidt RC, Bart HL, Nyingi WD. 2017. Multi-locus phylogeny reveals instances of mitochondrial introgression and unrecognized diversity in Kenyan barbs (Cyprininae: Smiliogastrini). Molecular Phylogenetics and Evolution 111:35–43.

Sexton JP, Hangartner SB, Hoffmann AA. 2014. Genetic isolation by environment or distance: which pattern of gene flow is most common? Evolution 68:1–15.

Simão FA, Waterhouse RM, Ioannidis P, Kriventseva EV, Zdobnov EM. 2015. BUSCO: assessing genome assembly and annotation completeness with single-copy orthologs. Bioinformatics 31:3210–3212.

Sloan DB, Havird JC, Sharbrough J. 2017. The on-again, off-again relationship between mitochondrial genomes and species boundaries. Molecular Ecology 26:2212–2236.

Struck TH, Cerca J. 2019. Cryptic species and their evolutionary significance. In: John Wiley & Sons L, editor. eLS. p. doi:10.1002/9780470015902.a9780470028292.

Thanou E, Kornilios P, Lymberakis P, Leaché AD. 2019. Genomic and mitochondrial evidence of ancient isolations and extreme introgression in the four-lined snake. Current Zoology 66:99–111.

Theim TJ, Shirk RY, Givnish TJ. 2014. Spatial genetic structure in four understory *Psychotria* species (Rubiaceae) and implications for tropical forest diversity. American Journal of Botany 101:1189–1199.

Therkildsen NO, Palumbi SR. 2016. Practical low-coverage genomewide sequencing of hundreds of individually barcoded samples for population and evolutionary genomics in nonmodel species. Molecular Ecology Resources 17:194–208.

Thomas L, Underwood JN, Adam AAS, Richards ZT, Dugal L, Miller KJ, Gilmour JP. 2020. Contrasting patterns of genetic connectivity in brooding and spawning corals across a remote atoll system in northwest Australia. Coral Reefs 39:55–60.

Thomaz D, Guiller A, Clarke BC. 1996. Extreme divergence of mitochondrial DNA within species of pulmonate land snails. Proceedings of the Royal Society of London. Series B: Biological Sciences 263:363–368.

Todaro MA, Fleeger JW, Hu YP, Hrincevich AW, Foltz DW. 1996. Are meiofaunal species cosmopolitan? Morphological and molecular analysis of *Xenotrichula intermedia* (Gastrotricha: Chaetonotida). Marine Biology 125:735–742.

Vieira FG, Lassalle F, Korneliussen TS, Fumagalli M. 2016. Improving the estimation of genetic distances from next-generation sequencing data. Biological Journal of the Linnean Society 117:139–149.

Villamor A, Costantini F, Abbiati M. 2014. Genetic structuring across marine biogeographic boundaries in rocky shore invertebrates. PLOS ONE 9:e101135.

Wang IJ, Bradburd GS. 2014. Isolation by environment. Molecular Ecology 23:5649–5662.

Westheide W, Schmidt H. 2003. Cosmopolitan versus cryptic meiofaunal polychaete species: an approach to a molecular taxonomy. Helgoland Marine Research 57:1–6.

Wickham H. 2016. ggplot2: Elegant Graphics for Data Analysis. New York: Springer-Verlag.

Worsaae K, Kerbl A, Vang Á, Gonzalez BC. 2019. Broad North Atlantic distribution of a meiobenthic annelid – against all odds. Scientific Reports 9:15497.

Woyke T, Teeling H, Ivanova NN, Huntemann M, Richter M, Gloeckner FO, Boffelli D, Anderson IJ, Barry KW, Shapiro HJ, et al. 2006. Symbiosis insights through metagenomic analysis of a microbial consortium. Nature 443:950–955.

Yeaman S, Whitlock MC. 2011. The genetic architechture of adatptation under migration-selection balance. Evolution 65:1897–1911.

Zakas C, Wares JP. 2012. Consequences of a poecilogonous life history for genetic structure in coastal populations of the polychaete Streblospio benedicti. Molecular Ecology 21:5447–5460.

