## Supplementary texts for "High genetic diversification in a symbiotic marine annelid is driven by microgeography and glaciation"

### Contents

Supplementary Text 1:

Bayesian models on co-regression between the linearised FST and geographic distance suggests a consistent isolation by distance pattern at the local and regional scales

Supplementary Text 2:

Implication of the consistent isolation by distance pattern in understanding population structuring among Elba populations indicates the sea level fluctuations during the last gracile cycles as major drivers

References for the Supplementary Texts

Supplementary Figure S1:

Cladogram of complete mitochondrial genomes with branch support

Supplementary Figure S2:

Animation of the 3D-MDS plot based on genetic distances for individuals from Elba locations color-coded by locations.

Supplementary Figure S3:

Animation of the 3D-MDS plot based on genetic distances for individuals from Elba locations color-coded by mitochondrial genome clades.

Supplementary Figure S4:

Workflow of reference transcript assembly.

Supplementary Figure S5:

Bayesian model-based analysis on the isolation by distance patterns between the local and regional scales.

**Supplementary Text 1:**

**Bayesian models on co-regression between the linearised $F_{ST}$ and geographic distance suggests a consistent isolation by distance pattern at the local and regional scales**

Bayesian models were applied to examine the IBD relationships at both local (Elba populations only) and regional (Elba and Mallorca populations) scales. Bayesian models were calculated using the R package *brms* v2.17.0 (Bürkner 2018), based on the generalised linear model without forcing zero-interval for robust testing with no assumptions regarding the origin. Only weakly informed priors were applied to the slope, intercept, and residual standard deviation sigma, upon sampling priors only from the model and testing against unfeasible priors restricting posteriors. Confirmation of model conversion and check against violation of assumptions were performed using the R packages *RStan* v2.26.13 (Stan_Development_Team 2021) and *DHARMa* v0.4.5 (Hartig 2022), respectively. Model representation was checked using the function *pp_check()* in *brms*. To model co-regression, first a Bayesian model was applied to one correlation (e.g., distance vs. $F_{ST}(1-F_{ST})$), then it was repeated after switching the predictor and response variables. Lastly the average slope was obtained from pairs of the two correlations in 2,000 draws each from the two model populations using the R package *posterior* v1.2.2 (Bürkner et al. 2022). Comparisons of correlation slopes and hypothesis testing were performed using the R packages *tidybayes* v3.0.2 (Kay 2019) and *bayestestR* v0.12.1 (Makowski et al. 2019).

The median slope for the regional scale encompassing Elba and Mallorca populations was slightly larger than the slope for the Elba local scale (Supplementary Figure S5; regional slope = 0.00175 vs. local slope = 0.00164). However, the posterior probability distribution of the slope difference showed that the posterior probability of the regional slope being larger than the local slope is only 74.7%. In other words, the posterior probability that the local-scale slope is larger than the regional-scale slope is 25.3%. Finally, the posterior probability that the difference of the slopes departs from zero by its 1 standard deviation is only 37.5%, and the probability is only 4.1% that the difference departs from zero by 2 standard deviation. These results therefore indicate that there is insufficient evidence to support that the slopes differ significantly from each other.

**Supplementary Text 2:**

**Implication of the consistent isolation by distance pattern points to the sea level fluctuations during the last gracile cycles as major drivers of population structuring around Elba populations**

Population structuring across Elba locations based on mitochondrial and exomewide SNP data showed that the limited dispersibility of *O. algarvensis* causes an effective reproductive isolation at the small spatial scales (<5 km). It can therefore be assumed that Elba populations and Mallorca populations are completely isolated today and that the genetic divergence between Elba and Mallorca populations is the function of time since an ancestral population diverged to derive today's Mallorca and Elba populations. The Mediterranean marine fauna vanished during the Messinian Salinity Crisis between 5.96 and 5.33 million years ago, until the Zanclean flood abruptly re-introduced seawater in the Mediterranean basin (Krijgsman et al. 1999; Harzhauser et al. 2007; Garcia-Castellanos et al. 2009). This point, 5.33 million years ago, was the possible earliest point in time when an ancestral population of *O. algarvensis* diverged to the present Mallorca and Elba populations, simply because no habitats around these islands have persisted from before this point. Given the consistent isolation by distance pattern over small and large spatial scales, if we assume that 5.33 million years ago as the timepoint when the split between Mallorca and Elba population occurred, we can estimate the timeframe of population genetic divergence around Elba populations. A correlation between the time and linearised $F_{ST}$ (i.e., $F_{ST}/(1-F_{ST})$) provides an estimation that population differentiation among Elba populations have developed since the 350,000 and 20,000 years ago, depending on the pair of locations. This timeframe estimate corroborates well with the population structure around Elba reflecting the shoreline structure during the Würm glaciation between ca. 115,000 and 11,700 years ago in the Late Pleistocene (Figure 5; Bossio et al. 2000). Furthermore, this estimate points towards the historical influence on the population structure during the Middle Pleistocene (781,000 ~ 126,000 years ago) before it led to further population structuring during the last glacial period in the Late Pleistocene.

Between 350,000 and 20,000 years ago, the sea levels fluctuated several times from the present level to 100 m levels below the present level (Hansen et al. 2013). Given the natural distribution of *O. algarvensis* in the proximity of seagrass meadows in shallow waters where light is available (Giere and Erséus 2002), the occurrence of the gutless oligochaete has likely been concentrated around historical distribution patterns of seagrass meadows. The horizontal distribution of seagrass meadows likely shifted according to the sea level fluctuations (Chefaoui et al. 2017). During the sea level declines, the seagrass meadows could extend towards offshore and thus the neighbouring seagrass habitats potentially merged or separated depending on the topology of the ocean floor. When the sea level was on the rise, the seagrass habitats could have reversed the movements. This fluctuation of sea level and corresponding shifts in positions of seagrass habitats during the Middle and Late Pleistocene give a probable explanation for the population differentiation across Elba locations.

### References for the Supplementary Texts


Bossio A, Cornamusini G, Ferrandini J, Ferrandini M, Maria Foresi L, Mazzanti R, Mazzei R, Salvatorini G, Sandrelli F. 2000. Dinamica dal Neogene al Quaternario della Corsica orientale e della Toscana. In. L’attività scientifica delle Università di Pisa e Corte. Pisa: Edizioni ETS. p. 87-95.

Bürkner P-C. 2018. Advanced Bayesian multilevel modeling with the R Package brms. The R Journal 10:395-411.

Bürkner P-C, Gabry J, Kay M, Vehtari A. 2022. posterior: Tools for Working with Posterior Distributions. https://mc-stan.org/posterior/.

Cerca J, Purschke G, Struck TH. 2018. Marine connectivity dynamics: clarifying cosmopolitan distributions of marine interstitial invertebrates and the meiofauna paradox. Marine Biology 165:123.

Chefaoui RM, Duarte CM, Serrão EA. 2017. Palaeoclimatic conditions in the Mediterranean explain genetic diversity of Posidonia oceanica seagrass meadows. Scientific Reports 7:2732.

Garcia-Castellanos D, Estrada F, Jiménez-Munt I, Gorini C, Fernàndez M, Vergés J, De Vicente R. 2009. Catastrophic flood of the Mediterranean after the Messinian salinity crisis. Nature 462:778-781.

Giere O, Erséus C. 2002. Taxonomy and new bacterial symbioses of gutless marine Tubificidae (Annelida, Oligochaeta) from the Island of Elba (Italy). Organisms Diversity & Evolution 2:289-297.

Hansen J, Sato M, Russell G, Kharecha P. 2013. Climate sensitivity, sea level and atmospheric carbon dioxide. Philosophical Transactions of the Royal Society A: Mathematical, Physical and Engineering Sciences 371.

Hartig F. 2022. DHARMa: Residual diagnostics for hierarchical (multi-level / mixed) regression models. https://CRAN.R-project.org/package=DHARMa.

Harzhauser M, Kroh A, Mandic O, Piller WE, Göhlich U, Reuter M, Berning B. 2007. Biogeographic responses to geodynamics: A key study all around the Oligo–Miocene Tethyan Seaway. Zoologischer Anzeiger - A Journal of Comparative Zoology 246:241-256.

Kamel SJ, Grosberg RK, Addison JA. 2014. Multiscale patterns of genetic structure in a marine snail (*Solenosteira macrospira*) without pelagic dispersal. Marine Biology 161:1603-1614.

Kay M. 2019. tidybayes: Tidy data and geoms for Bayesian models. http://mjskay.github.io/tidybayes/.

Kelly RP, Palumbi SR. 2010. Genetic Structure Among 50 Species of the Northeastern Pacific Rocky Intertidal Community. PLOS ONE 5:e8594.

Krijgsman W, Hilgen FJ, Raffi I, Sierro FJ, Wilson DS. 1999. Chronology, causes and progression of the Messinian salinity crisis. Nature 400:652-655.

Makowski D, Ben-Shachar MS, Lüdecke D. 2019. bayestestR: Describing effects and their uncertainty, existence and significance within the Bayesian framework. Journal of Open Source Software 4:1541.

Stan_Development_Team. 2021. RStan: the R interface to Stan. https://mc-stan.org/.

Villamor A, Costantini F, Abbiati M. 2014. Genetic structuring across marine biogeographic boundaries in rocky shore invertebrates. PLOS ONE 9:e101135.
