## Supplemental Figure S1 for "High genetic diversification in a symbiotic marine annelid is driven by microgeography and glaciation"

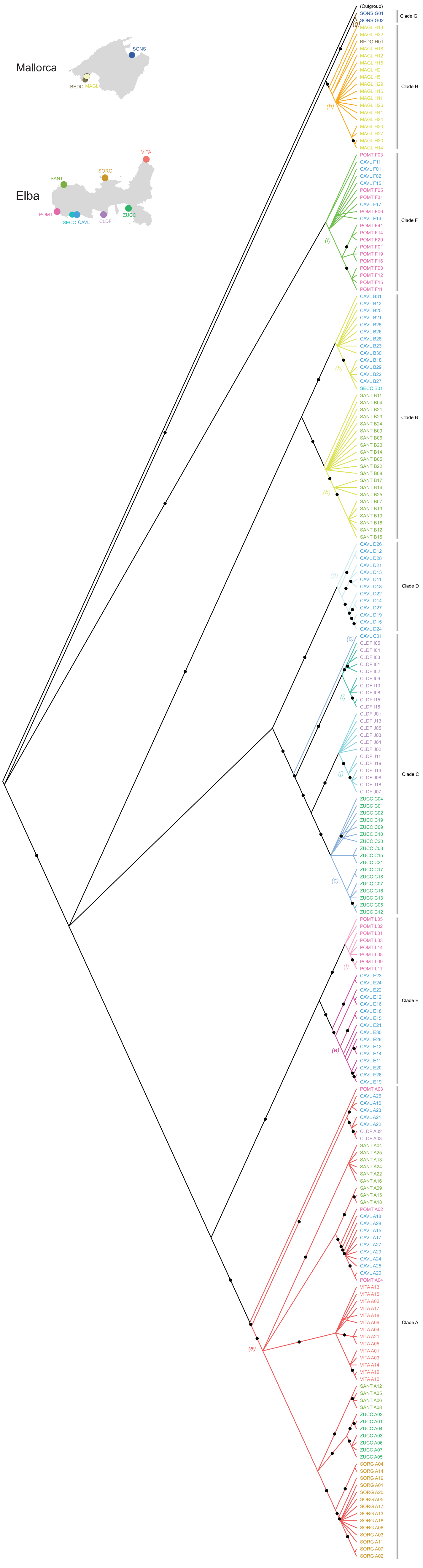


**Supplementary Figure S1 Cladogram of complete mitochondrial genomes with branch support**. The colors of terminal labels correspond to their locations indicated on the maps. The colors of branches and small italic letters in brackets indicate haplogroups based on COI sequences. Major clades based on the complete mitochondrial genomes are indicated on the right. Posterior probability of clades >0.95 are shown with black circles. Phylogeny was calculated using the same procedure as for Figure 3.
