## Supplementary figures and images for "High genetic diversification in a symbiotic marine annelid is driven by microgeography and glaciation"

### Supplemental Figure S2

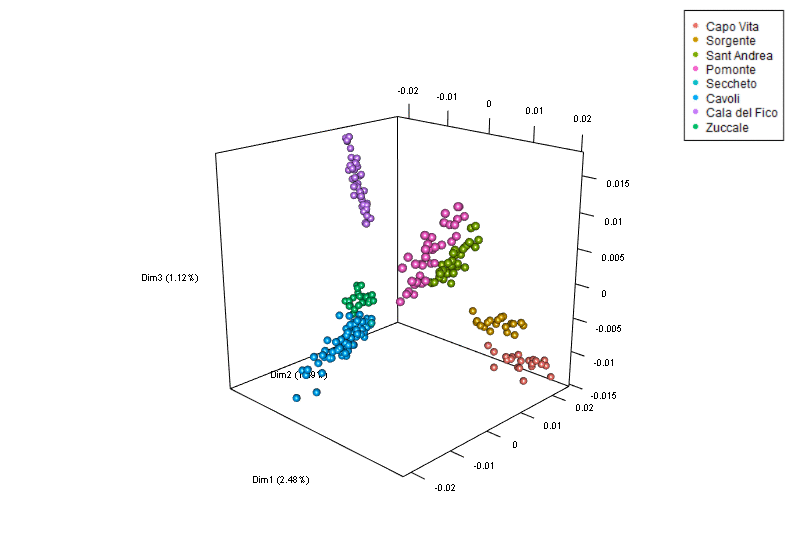

### Supplemental Figure S3

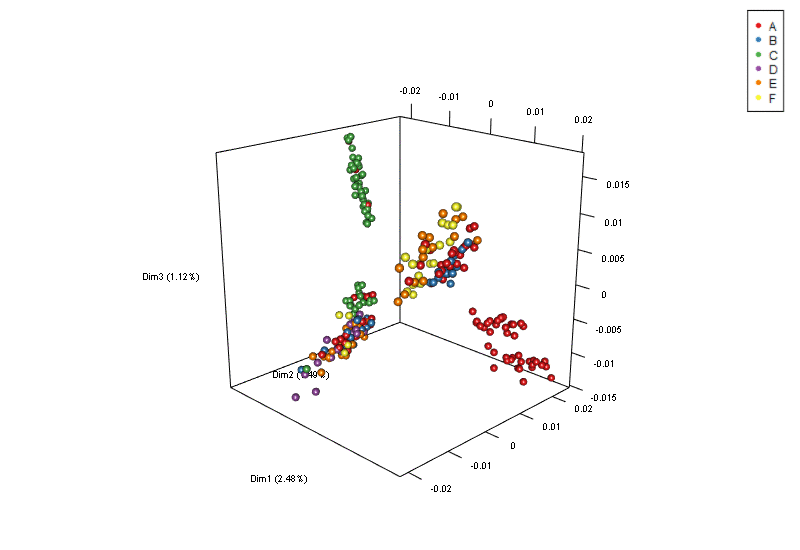
