## Supplemental Figure S4 for "High genetic diversification in a symbiotic marine annelid is driven by microgeography and glaciation"

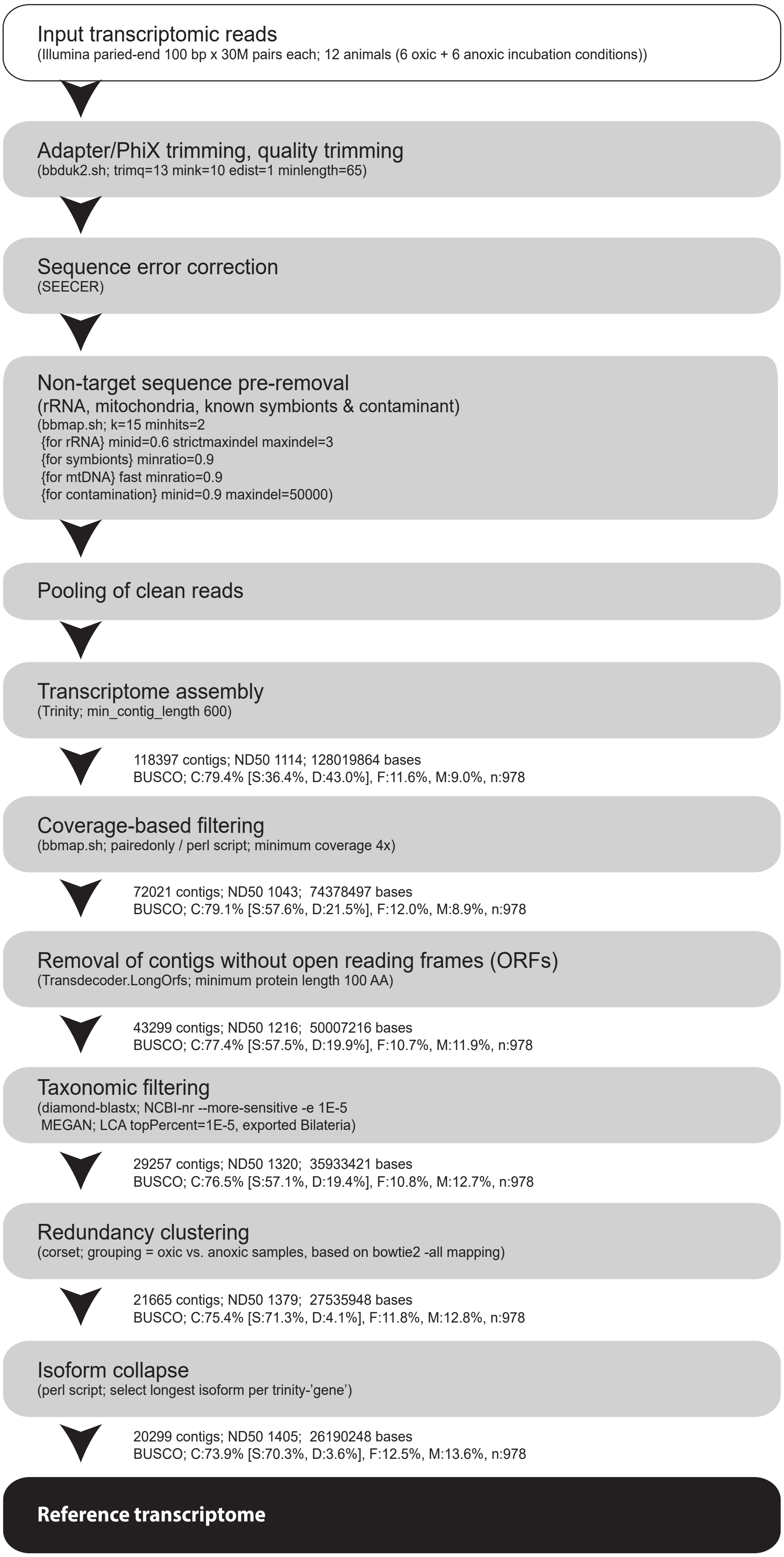


**Supplementary Figure S4 Workflow of reference transcript assembly**. Each box indicates the objective in the workflow, software implemented, and parameter setting that differ from the default setting. Statistics indicated along with the arrow show the assembly information including the total number of contigs, the nucleotide length of ND50 contig (50%tile according to the contig length), total number of nucleotides, and the BUSCO completeness assay results (C; complete, S; single genes, D; duplicated genes, F; fragmented genes, M; missing genes, and n; the total number of genes assessed.
