## Supplemental Figure S5 for "High genetic diversification in a symbiotic marine annelid is driven by microgeography and glaciation"

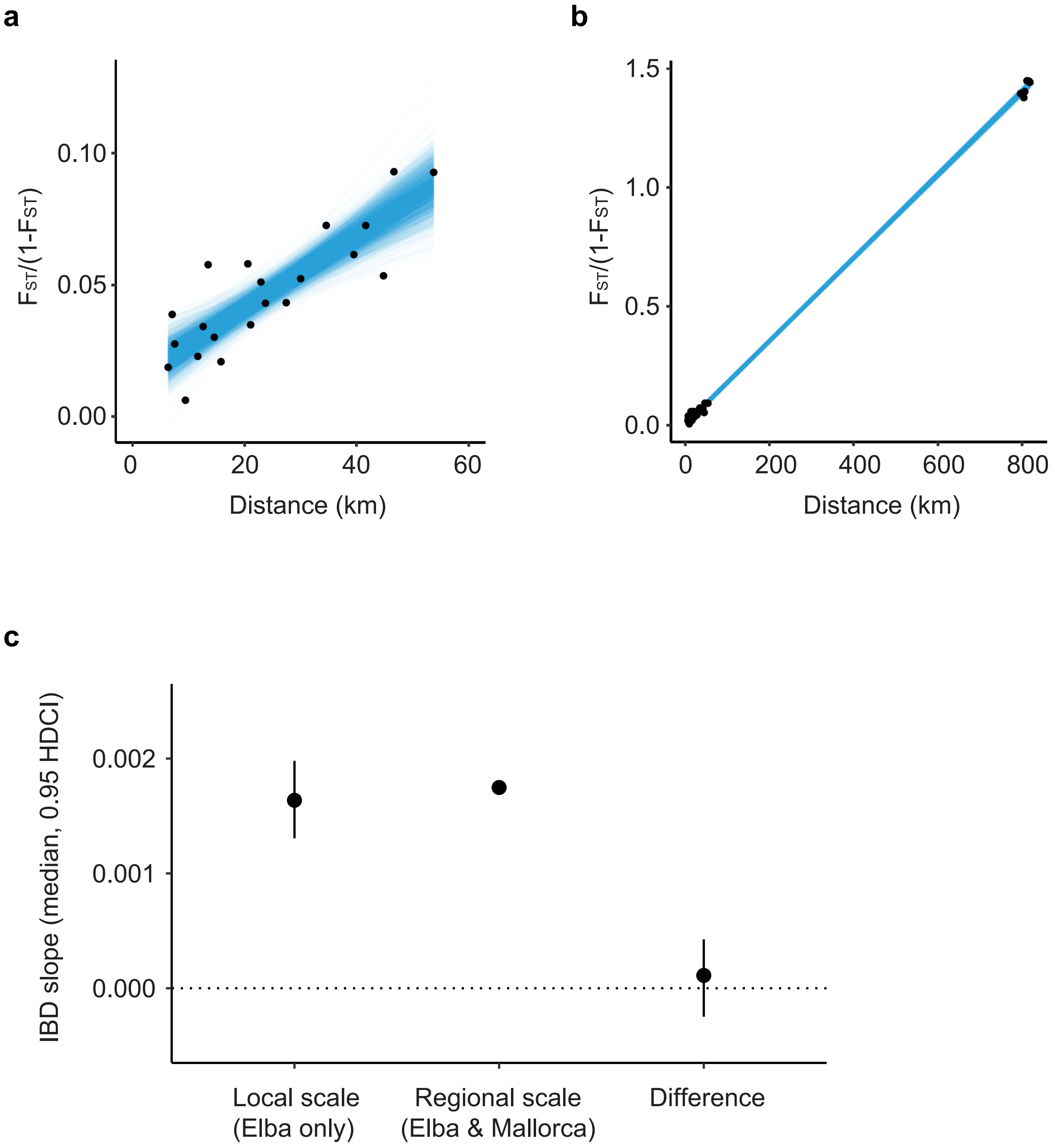


**Supplementary Figure S5 Bayesian model-based analysis on the isolation by distance patterns between the local and regional scales**. (**a, b**) Summary plots of the distance-linearised $F_{ST}$ correlations at the local scale (a; Elba only) and regional scale (b; Mallorca and Elba populations). Black dots show observed data, and each blue line represents a posterior coregression slope out of 2,000 random draws from each of the Bayesian populations. (**c**) Distributions of co-regression slopes for the local and regional scales and the difference between the two slopes, based on 2,000 random pairs drawn from the Bayesian populations. The circles and vertical bars indicate the median and 95% highest density credible intervals, respectively.
